# Multi-scale Engineered Vasculature and Hierarchical Porosity via Volumetric Bioprinting-guided Photopolymerization-induced Phase Separation

**DOI:** 10.1101/2025.07.21.665988

**Authors:** Oksana Y. Dudaryeva, Maj-Britt Buchholz, Gabriel Größbacher, Sofia Amaral, Sammy Florczak, Alvaro Rojo Ferrer, Mark W. Tibbitt, Riccardo Levato

## Abstract

Vascularization remains a major challenge in hydrogel-based engineered tissues due to the inherent nano-scale porosity of common synthetic and natural biomaterials. Critically, the confinement imposed by the nanoscale network inhibits the outgrowth of blood vessels required for oxygen and nutrient delivery. Despite advancements in biofabrication that enable formation of small channels (0.1–1 mm), achieving vascularization (with capillaries down to 10 µm) throughout cm-scale bioprinted constructs remains a critical bottleneck. Herein, we integrated phase separating cell–interactive gelatin–norbornene hydrogels with volumetric bioprinting to generate architecturally defined centimeter-scale constructs with 0.1–1 mm scale printed channels and interpenetrating micron-scale porosity. Our novel approach allowed us to generate freeform construct designs with light-controllable micron-scale and hierarchical porosity. Importantly, this porosity enabled endothelial cell infiltration and microvessel outgrowth deep into the engineered tissue. Micron-scale vascular structures formed in the pore spaces with feature sizes on the scale of capillaries (<10 µm), crucial to provide oxygen and nutrients to all regions of the hydrogel. The networks remained stable for over 14 days, outperforming classical nanoporous biomaterials. These complex hydrogel-based constructs with engineered multi-scale vascular networks have substantial potential for the generation of actively perfusable advanced tissue models.

## Introduction

Blood vessels form complex networks that interpenetrate and perfuse tissues and organs, providing critical nutrients and oxygen while removing metabolic waste. In addition, vascular networks are dynamic and rapidly adapt to physiologic changes to meet the changing metabolic demands of tissues. In this context, they are essential for maintaining tissue homeostasis and supporting cellular function. Vascular disorders, such as atherosclerotic cardiovascular disease, that impair this crucial function remain the leading cause of mortality across all developed societies and exacerbate comorbidities, including hypertension, hyperlipidemia, and pain disorders.^1^ Restriction of blood supply as a consequence of vascular disorders leads to ischaemia, nutrient depletion, and waste accumulation, ultimately causing tissue or organ necrosis or death.

Engineering vasculature in vitro is therefore essential for biological investigation of vascular function and for the fabrication of functional tissue replacement for regenerative medicine. However, the creation of large, fully vascularized, clinically relevant constructs remains a major challenge. Several strategies have been explored to produce viable and functioning vasculature in vitro in 3-dimensional (3D) engineered tissues. Often hydrogels based on natural ECM components, such as collagen, or fibrin are used to generate in vitro vasculature with limited control of the vascular architecture.^2,3^ To generate topologically defined vascular features within natural hydrogels, microfabrication or advanced sacrificial molding strategies have been applied.^4,5^ While effective at small scales, this strategy is limited mainly to fabrication of enclosed microfluidic chips, which are necessary to maintain the mechanical integrity of the natural hydrogels during long-term culture. The difficulties in handling these soft hydrogels and their limited stability impede their use in the broader context of biofabrication, especially for producing large-scale constructs with complex shapes and architectural cues, i.e. tissue-specific geometries with multi-scale branching or interconnected loops that mimic native vasculature.

In this context, the ability to generate implantable vascularized hydrogel constructs with appropriate stability and structural integrity while capturing shape-complexity is desirable. Photocurable synthetic hydrogels are often used in biofabrication as they enable tailored structural cues due to their spatioselective photocrosslinking and offer appropriate stability and structural integrity. However, synthetic photocrosslinkable 3D biomaterials come with their own shortcomings due to their inherent nanoporosity, stiffness, or limited viscoelasticity that often hinders effective outgrowth of 3D vascular structures.^6,7^ To overcome this limitation, light-based bioprinting techniques, including digital light processing (DLP) and volumetric bioprinting (VBP),^8,9^ have emerged as effective methods to create intricate networks of 3D channels using photoresponsive hydrogels that recapitulate the complex anatomical architecture of native vasculature.^10,11,12^ These permissive channels can be populated with living cells outgrowing into vascular networks accelerating progress toward the in vitro fabrication of large-scale vasculature. The bioprinting techniques, such as DLP and VBP, can effectively pattern large channels to establish static vessels in the 0.05–1 mm range, recapitulating the hierarchical and relatively static architecture of large-scale vasculature. The dimensions of the microvasculature (< 10 μm) are more challenging to capture with these techniques especially in the context of their randomly organized, highly dynamic nature that is capable of continuous sprouting and remodeling in response to external cues.^11,13^ One strategy that overcomes the resolution limit is the use of multiphoton lithography, which enables sub-micron feature generation but requires impractical amounts of time to construct large, clinically-relevant microvascular networks, accommodating their dynamic remodelling.^14,15,16^

Therefore, advanced biofabrication techniques that facilitate rapid formation of dense networks of capillaries less than 10 µm in size are needed to enable nutrient exchange across large volume engineered tissue models or to avoid rejection of biomaterial-based implants. Ideally, such approaches would also facilitate the interface between channels and vessels (0.1–1 mm) and the generated microvasculature, enabling facile perfusion of the entire construct. Such multi-scale engineered vascular networks would address the open challenge of microvascularization of in vitro tissue models, which is a determining factor for their success in disease modeling and drug screening pipelines.

In this work, we generated bioprinted constructs from materials that enable vascularization of cm-scale in vitro tissue models. We fabricated materials permissive for cell infiltration and vasculature outgrowth using photocontrolled phase-separating bioinks based on the commonly used biomaterials: gelatin and poly(ethylene glycol) (PEG). Here, the integration of gelatin is particularly advantageous and cost-effective for bioprinting applications due to its widespread availability, intrinsic bioactivity, and low material cost. The presence of cell-adhesive motifs (RGD sequences) and enzymatically degradable sites [matrix metalloproteinase (MMP)-sensitive domains] makes gelatin highly suitable for large-scale cell-based applications, eliminating the need for challenging and expensive chemical modifications like peptide conjugation. Utilization of these materials for volumetric bioprinting enabled generation of bioprinted constructs with multi-modal porosities of a controllable size. Importantly, this enabled us to generate prints with large pores or channels (mm-scale) interconnected with micron-sized channels (< 10 μm) within a single porous construct. To our knowledge, this is one of the few strategies that are capable of providing control across multiple scales: shape architecture control at the millimeter-scale combined with control over the intrinsic porosity at micron-scale. These multi-scale constructs were populated with vascular endothelial cells to create endothelial structures down to the microcapillary range. Generation of microvasculature in vitro using macroporous hydrogels will expedite the production of large vascularized in vitro models with tunable architectures. Such models will likely enable controlled parametric studies that will fundamentally enhance our understanding of angiogenesis in the course of disorders, and accelerate production of vascularized engineered tissues and organs.

## Results and Discussion

### Engineering phase-separating bioinks for volumetric bioprinting

Previously, we and others have established macroporous hydrogel biomaterials through photopolymerization-induced phase separation.^17–23^ In this process, an initial single-phase aqueous polymer solution phase separates during the polymerization process, enabling the formation of an interconnected network of micron-scale pores under suitable conditions (**Figure 1a**).^17^ We formulated phase separating bioresins based on gelatin and PEG polymers with controllable interconnected porosity based on thiol–ene chemistry to facilitate photopolymerization-induced phase separation during crosslinking. The formulation consisted of gel forming components [norbornene-functionalized gelatin (GelNb) and thiol-functionalized 4-arm poly(ethylene glycol) (PEG-SH; 10kDa)]; an excluding phase [dextran (MW ≈ 100 kDa)]; and a viscosity modifier [hyaluronic acid (HA; *M*_n_ ≈ 2 MDa)] (**Figure 1b**).

**Figure 1.**
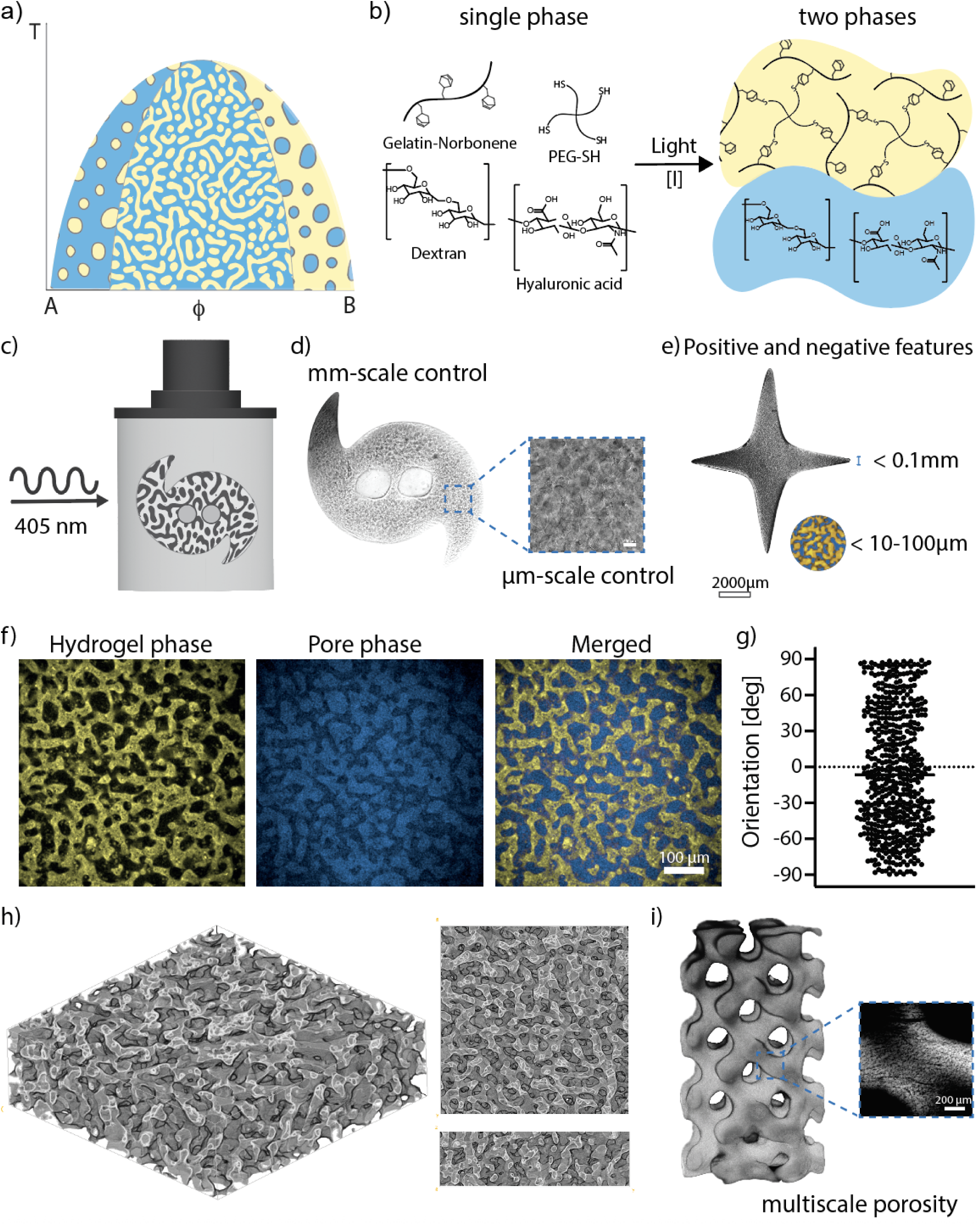
Formation of porous hydrogels from gelatin and PEG, and polysaccharides. a) In a multi-component system the initially miscible macromolecules (gelatin and polysaccharides) become immiscible upo photocrosslinking of one of the components. Growth of the molecular weight induces phase separation and pore formation, either through binodal (formation of spheres) or spinodal decomposition mechanisms. b) Our phase-separated system is based on photocrosslinkable GelNb–PEG-SH macromolecules and an excluded phase containing the polysaccharides dextran and HA. c) The phase-separating materials can be volumetrically printed generating complex 3D objects with intrinsic interconnected porosity. d) Printing macroporous hydrogels provides control over mm- and micron-scale features. Transmitted light image. e) The large-scale features can be printed with a submillimeter resolution, the microscale features are printed with varying resolution in the range of 5-100 μm. Cropped transmitted light image. f) Photocrosslinking of Gelatin-NB/PEG-SH (gel-phase; yellow) in presence of dextran/HA (excluded phase; blue) crosslinked at 5.6 mW/cm^2^ yielded hydrogel materials with interconnected perfusable porosity, demonstrated by perfusion of the pore phase with high Mw dextran-FITC (Mw=500 kDa). g) The spinodal porosity is isotropic in all directions as shown by the quantitative analysis of the pore orientation. h) Three-dimensional reconstruction of the phase-separated hydrogel reveals a continuous porous network throughout the volume. The image was reconstructed from confocal z-stacks and rendered using the Volume Viewer plugin in Fiji. i) Multiscale porous gyroid architectures were printed at *I*_av_ = 6 mW/cm^2^ featuring designed macroporosity (∼1 mm) and intrinsic, phase separation-derived microporosity (13.5 ± 2.9 µm). Scale bar = 200 μm.

In regenerative medicine and tissue engineering, there remains a major challenge in capturing the hierarchical architecture of living tissues, which spans from subcellular-sized features to large anatomical structures. In this work, we sought to apply this concept in the design of biofunctional macroporous bioinks suitable for volumetric bioprinting (VBP) to engineer constructs with controlled millimeter-scale features and micron-scale interconnected porosity (**Figure 1c,d**). This allowed us to generate structures via VBP with features with size below 0.1 mm and micron scale porosity in the ranges of 5–100 μm. Crosslinking of GelNb–PEG-SH hydrogels in the presence of dextran and HA generated materials with interconnected and perfusable porosity as demonstrated by perfusion of the porous space with large molecular weight Dextran-FITC (500 kDa; **Figure 1f**). Generating well-defined porosity in gelatin-based systems was challenging due to temperature-dependent physical interactions between gelatin macromolecules. At room temperature gelatin chains exhibit physical interactions leading to uncontrolled aggregation of the chains which could trigger phase separation before polymerization and affect the architecture and pore size of the final structures (**Figure S1**). Maintaining a constant temperature was important for reproducible results. We maintained the temperatures above 37 °C (38 – 45°C) during crosslinking to yield bicontinuous network structures formed via spinodal decomposition. The porosity was isotropically distributed throughout the hydrogel volume as shown by porosity orientation (**Figure 1g,h**). The phase-separated inks were used to generate constructs with multi-scale porosity, such as a gyroid that was designed with large-scale printed porosity of 1 mm and exhibited small-scale macroporosity of 13.5 ± 2.9 μm pores driven through phase-separation which can potentially enable enhanced mass transport and cell infiltration (**Figure 1i**).

### Controlling porosity with formulation, irradiation intensity, and volumetric light fields

To understand the design parameters for photopolymerization-induced phase separation in this system, we first assessed how the formulation of the phase-separating inks affected the final material porosity in gels prepared by casting the polymer solution into molds, followed by crosslinking at a uniform cytocompatible polymerization intensity (I = 5 mW/cm^2^; λ = 405 nm). The final material porosity depended on the concentration of both dextran and HA. The size of the final material porosity scaled proportionally with dextran concentration from 8.7 ± 2.2 to 19.4 ± 3.9 μm, at 0 and 5 wt% dextran respectively (**Figure S2**) and decreased with an increasing concentration of HA from 23.2 ± 1.6 to 7.2 ± 1.3 μm at 0.05 wt% and to 0.3 wt% HA (**Figure S3**). We next demonstrated that we could spatially control pore size by modulating the polymerization kinetics. Given that kinetics can be governed by the irradiation conditions for thiol–ene photopolymerizations, this was achieved by varying the irradiation intensity without changing dextran or HA concentration (**Figure 2a**).^18,19,22^ Increasing the irradiation intensity decreased the time to reach complete gelation in our GelNb–PEG-SH based bioinks, which was indicated by the time to reach plateau storage modulus (G’, **Figure S4**). The polymerization kinetics determine the time to reach network percolation, which competes with the dynamics of the phase separation process. Extending the time until full percolation allowed the system to proceed further in the phase separation, leading to larger pore sizes and providing a handle to tune the final porosity in the material from a single formulation. We generated different porosities from a single formulation by polymerizing the hydrogels using different irradiation intensities under LED light (**Figure 2b)**. We used the BoneJ plugin in Fiji to quantify the porosity in both 2D and 3D that featured pore segmentation, skeletonization, and determination of the pore widths (**Figure 2c,d**).^24,25^ Decreasing the irradiation intensity from 10 to 0.5 mW/cm^2^ generated materials with pores ranging from 13.0 ± 2.4 to 52.9 ± 18.2 µm, respectively (**Figure 2e**). While the porosity fraction remained largely consistent across the range of polymerization intensities, a minor reduction was noted at the highest intensity (10 mW/cm²) that was significant in comparison with the medium polymerization intensity (5.6 mW/cm²). The fraction of porosity was 0,38 ± 0,06 for 10 mW/cm² and 0,51 ± 0,04 for 5.6 mW/cm² intensities (**Figure 2f**).

**Figure 2.**
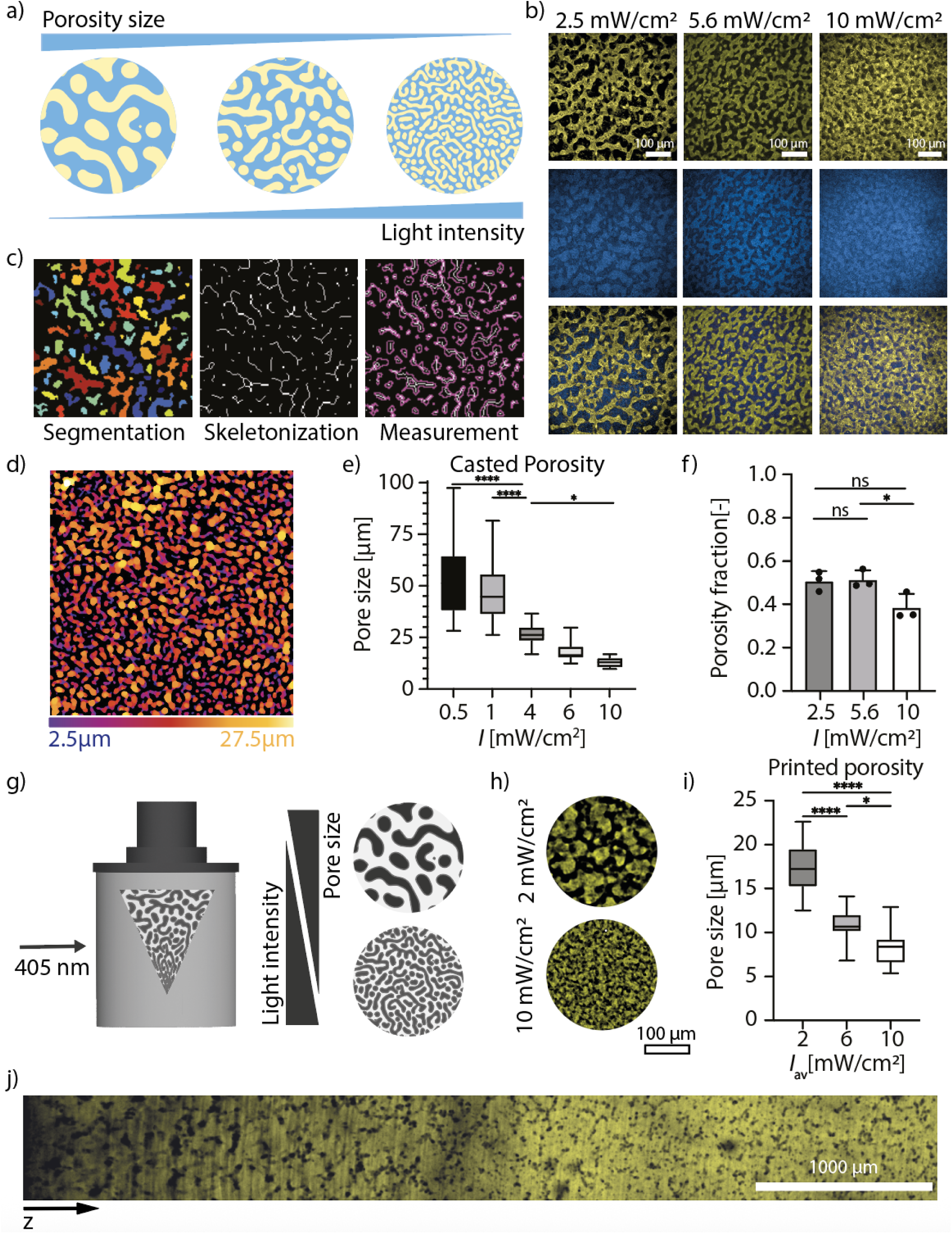
Controlling material porosity using light intensity. a) The size of porous features in photopolymerizable gelatin/PEG systems can be controlled using irradiation intensity, which governs polymerization kinetics. The final porosity increases at decreasing polymerization intensity. b) Confocal images depicting the porosity of the casted hydrogels, with pore size decreasing with increasing intensity. c) Representation of the porosity analysis pipeline. In order to characterize the pore sizes of the phase-separated systems the 3D micrograph stack was first segmented within ImageJ and then processed in using the BoneJ plugin. This also enabled for a skeletonized depiction of the voids within the construct. d) The image presents a color-coded map of pore size distribution within the hydrogel matrix, where each color corresponds to a specific pore width. The porosity ranged between 2.5 and 27.5 μm for hydrogels polymerized at 2.5 mW/cm^2^. e) The porosity within the casted systems has been controlled between 13.0 ± 2.4 to 52.9 ± 18.2 µm for 0.5 and 10 mW/cm^2^ respectively (n = 3). Box plots represent the interquartile range (25th–75th percentile), medians, and whiskers show the full data range (minimum to maximum). Pore size decreases significantly with increasing light intensity (* = p < 0.05, **** = p < 0.0001; one-way ANOVA test). f) The fraction of porosity varied between 0.38 ± 0.06 for 10 mW/cm² and 0.51 ± 0.04 for 5.6 mW/cm² polymerization intensities (n = 3). Values are reported as mean and std. dev. (* = p < 0.05, ns = not significant). g) Volumetric printing enables generation of objects with varying porosity or with porosity gradients using volumetric light fields, either through printing objects at varying light intensity or applying the light fields with gradient intensity. h) The porosity of the printed objects was controlled by printing objects at different light intensities. i) The porosity was controlled between 17.5 ± 3.2 μm at *I_av_* = 2 mW/cm^2^ and 8.3 ± 2.0 μm at *I_av_* = 10 mW/cm^2^ respectively (n = 3). Box plots represent the interquartile range (25th–75th percentile), medians, and whiskers show the full data range (minimum to maximum). (* = p < 0.05, **** = p < 0.0001; one-way ANOVA test). j) A cylindrical object was printed with the light field with gradient *I_av_*decreasing along the cylinder’s z-axis from 8 to 0.8 mW/cm^2^, generating porosity gradient with the porosity sizes increasing along the decreasing *I_av_*. The tile scan image was taken along the z-axis of the cylinder. Scale bar = 1000 μm.

These unique properties of the phase-separating bioresins enable new directions in their use in VBP, which operates with locally controllable light fields, wherein phase separation can be used to form objects with locally controlled micron-scale porosity. In this manner, we could form objects with different intrinsic porosity through printing objects at different light intensity or objects with porosity gradients, by applying the light fields with gradient intensity (**Figure 2g**). We demonstrated this by volumetrically printing objects at different irradiation intensities (**Figure 2h**). We printed the objects with intrinsic porosities ranging between 17.5 ± 3.2 μm and 8.3 ± 2.0 μm in a single print volume, polymerizing the objects at different average irradiation intensities (*I*_av_) of 2 and 10 mW/cm^2^, respectively (**Figure 2i**). The difference between casted and printed porosity might stem from the light intensity distribution across the print volume during the volumetric printing and potential differences in oxygen concentrations during crosslinking.

Although there were differences in pore sizes between the casted gels and printed gels at equivalent irradiation intensities, the scaling of pore size with intensity was conserved. While the process allows for control over the average irradiation intensity across the construct, localized regions can experience peak intensities up to 4–8 times higher than the average (depending on settings and the printed architecture), potentially influencing phase separation dynamics and pore formation (**Figure S5**). We further used a custom-built VBP module to generate a hierarchical porosity within a single construct. By applying a volumetric light projections with gradient irradiation intensity linearly decreasing from *I_av_* = 8 to 0.8 mW/cm^2^ along the z-axis we printed a cylindrical construct with a continuous porosity gradient increasing from 14.7 ± 1.6 μm to 67.7 ± 12.5 μm (**Figure 2j**).

### Vascularization of macroporous printed constructs

The porosity of phase separating hydrogels has been shown to be permissive for the cell infiltration, migration, and outgrowth of multicellular structures.^18,19^ We hypothesized that the interconnected pore space within our hydrogels would be permissive for the outgrowth of the vascular structures potentially vascularizing volumetrically printed constructs with different sizes and architectures. To enable the formation of microcapillary-like structures with diameters below 10 μm within volumetrically printed constructs, we precisely modulated the printing intensity (*I_av_*= 8–10 mW/cm²) to control phase separation dynamics during printing. Using this approach we fabricated constructs with intrinsic porosity below 10 μm, that would limit the outgrowth of larger vessels and favor development of microvascular networks within the printed architectures. To demonstrate pore permissiveness at scale relevant for mammalian cells we infused the pore space of the printed hydrogels with green fluorescent microbeads (d = 6 μm) that infiltrated in the pores of the construct (**Figure S6**).

To generate vasculature within the macroporous space of our printed constructs, we seeded GFP-expressing human umbilical vein endothelial cells (HUVECs-GFP) in combination with human mesenchymal stem cells (hMSCs) on top of the printed structures (**Figure 3a**). The seeded cells cooperatively evolved into vascular-like networks on the surface of the printed constructs and within the pore space with a mean diameter of 13.2 ± 5.7 μm over the period of 3 to 5 days.

**Figure 3.**
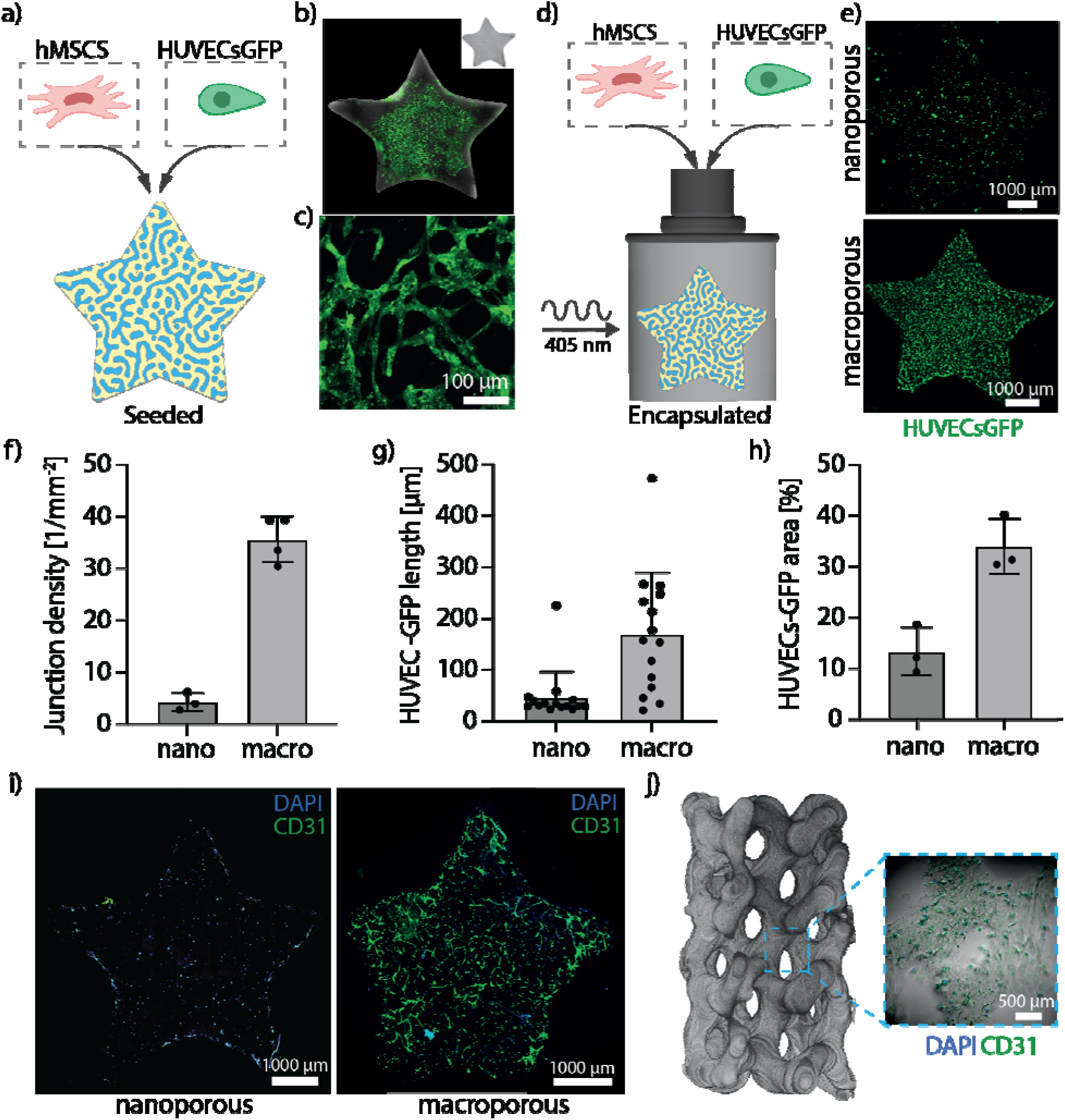
Macroporosity enhances vascular network formation in bioprinted constructs. a) The volumetrically printed shapes were seeded with the HUVECs-GFP and hMSC. b) The formed endothelial network partially vascularized the porous space of the printed constructs showing limited ingrowth into material bulk. b) Endothelial structures in the porous space of the seeded hydrogel constructs. c) To improve vascularization of the printed construct the phase-separating gels were volumetrically printed with cells suspended in gel forming solution. d-e) The printed constructs with nano- and macroporosity exhibited different cell infiltration and outgrowth of HUVECs-GFP. f) The total junction density was 4.3 ± 1.8 mm^-2^ in nano- and 37.4 ± 3.4 mm^-2^ in macroporous constructs (N = 3). g) The average vessel length was 33.6 ± 3.0 in nanoporous and 169.7 ± 30.9 μm in macroporous printed constructs (N = 3). h) The total cell area was 13.4 ± 4.8 in nano and 33.99 ± 5.46 % in macroporous constructs (N = 3). All values in f-h are reported as mean and std. dev. i) Confocal tile scan of cell-laden bioprinted stars with nano- and macroporosity. In nanoporous prints HUVECs-GFP appeared mostly rounded. HUVECs-GFP organized int the CD31 positive nascent vascular structures within the macroporous construct with an average diameter of 8.2 ± 2.7 μm. j) Light-sheet reconstruction of the gyroid with multi-scale porosity laden with hMSCs and HUVECs-GFP (1:1, 500*10^3^ cells/ml), that developed vascular structures in the macroporous space (cropped image).

These endothelial networks grew into the pore space to depths of ≈ 300 μm but did not populate the whole volume of the prints, partially due to an uneven distribution of the cells on the surface of the constructs during seeding (**Figure 3b,c; Figure S7**). To improve population of the porous space, we volumetrically printed the constructs with cells suspended in the gel forming bioink (*I_av_* = 8–10 mW/cm²) evenly distributing the cells in the in situ phase separating gels (**Figure 3d**). We fabricated the constructs both with nano- and macroporosity to assess if micron-scale porosity improves vascular outgrowth in 3D space compared to the nanoporous hydrogels. Cell viability assessed 24 h after printing was comparable between the two groups, 73.8 ± 5.7% in nanoporous hydrogels and 71.3 ± 17.9% in macroporous hydrogels (**Figure S8**).

After 5 days of culture, we analyzed the morphology and outgrowth of HUVECs-GFP within nano- and macroporous hydrogels. GFP-positive cells within the nanoporous hydrogels were rounded, while macroporous prints exhibited more spreading and organization (**Figure 3e**). The endothelial networks developed throughout the entire volume of the printed constructs (**Figure S9**).

The average vessel length was 33.6 ± 3.0 in nanoporous and 169.7 ± 30.9 μm in macroporous printed constructs (**Figure 3g**). The total cell area percentage taken up by HUVECs-GFP was 13.4 ± 4.8% in nano- and 33.9 ± 5.5% in macroporous constructs (**Figure 3h**). The total junction density of vascular structures was 4.3 ± 1.8 mm^-2^ in nano- and 37.4 ± 3.4 mm^-2^ in macroporous constructs (**Figure 3f**). The nanoporous cell-laden prints exhibited limited to no cell spreading, save for the cells attached to the outer rim of the printed construct, while endothelial structures positive for CD31 have formed over a period of 5 days in porous constructs (**Figure 3i**). The endothelial structures within macroporous stars had an average vessel diameter of 8.2 ± 2.7 μm. Interestingly, over time some of the cell-laden macroporous prints exhibited shrinking while retaining the formed vascular structures (**Figure S10**). We also demonstrated formation of vascular structures in the larger constructs with multi-scale porosity by printing large gyroids with cells (**Figure 3j**), to underline the versatility of printing complex architectures in which cells can grow.

### Vascularization of perfused bioprinted chips

Finally, we generated perfusable systems to test if perfusion enhances the vascular outgrowth and increases the stability of the formed vascular structures. These systems hold potential to be used in combination with other cell types, spheroids or organoids to generate perfusable vascularized organ or tumor models. Earlier, researchers have demonstrated guidance of angiogenesis from larger vessels using cell-instructive cues, including growth factors, and mechanical stimulation using fluid flow.^26,27,28^ However, achieving vascularization of large perfusable constructs based on synthetic hydrogels remains a significant and unresolved challenge.

We volumetrically printed constructs with a large perfusable central channel (d ≈ 800 μm) both with nano- and macroporosity (**Figure 4a**). In constructs printed with phase-separating bioinks, interconnected porosity extended into the bulk of the material from the central channel. As a control, nanoporous constructs were printed using GelNb–PEG-SH without the excluding components, dextran and HA (**Figure 4b**), although the material bulk was nanoporous, the orthogonal striations have formed along the main channel, a common artefact of the volumetric printing (**Figure S10**). The channel of the constructs, designed with a u-shaped curvature to promote retention of the seeded cells, was seeded with the HUVECs-GFP and hMSCs (1:1 ratio; 10 μL, total of 1 x 10^6^ cells). Following seeding, the chips were rotated around their longitudinal axis for 6 h using a custom-built rotating device to ensure uniform distribution of cells along the channel walls. The seeding was repeated twice over a one-week pre-culture period.

**Figure 4.**
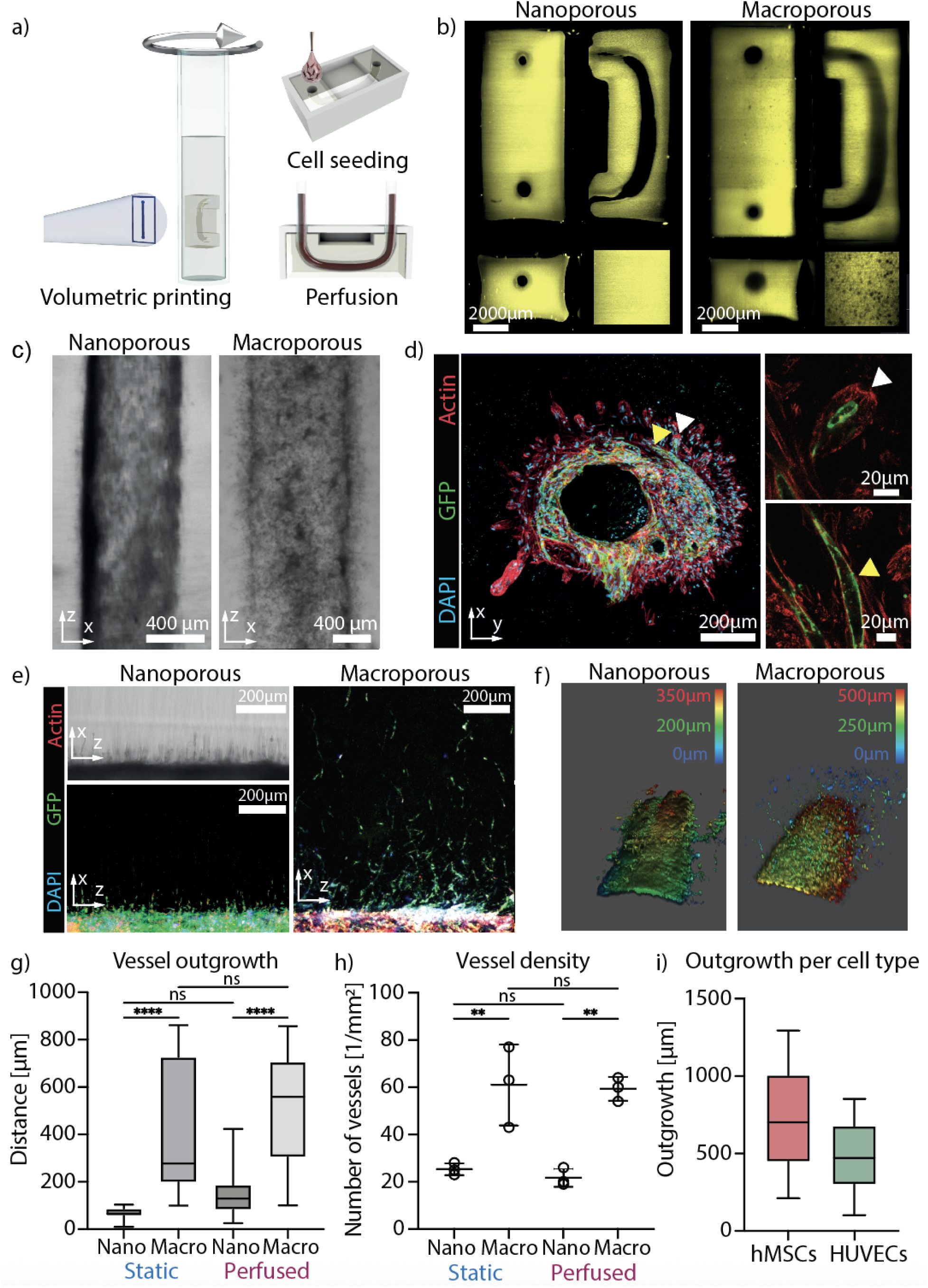
Vascularization of the printed perfusable constructs. a) Perfusable constructs volumetrically printed, with a central perfusable channel that was seeded with the hMSCs and HUVECs. b) Light-sheet reconstruction of the perfusable chips generated with nano- and macroporosity. c) After one week of pre-culture the cells in the nanoporous chips have shown limited outgrowth into the bulk of the construct; the macroporous chips exhibited improved cell infiltration from the channel. d) Left: The seeded hMSCs and HUVECs infiltrated the bulk of the constructs from the main channel, the hMSCs were followed by HUVECs; Top-right, bottom-right: hMSC-driven protrusions that are lined with HUVECs and lumenized. e) The macroporous chips (right) show enhanced vascularization compared to the chips with nanoporosity (left) along the length of the perfusable channel. The outgrowth in the nanoporous constructs is mainly realized through printing striations as shown by transmitted light micrograph (top left). f) 3D reconstruction of confocal z-stacks (500 μm) of the main channel in the nano- and macroporous perfused constructs showing enhanced outgrowth of the cells in macroporosity. g) Vascular outgrowth within nano- and macroporous constructs (N = 3). h) The vessel density in nano and macroporous constructs kept under static conditions was 25.3 ± 2.5 mm^-2^ and 61.0 ± 17.09 mm^-2^ respectively. In perfused constructs the vessel density was 21.7 ± 2.8 mm^-2^ and 59.3 ± 5.0 mm^-2^ for the nano- and macroporous construct respectively (N = 3). i) Analysis of cell-specific outgrowth showed that on average hMSCs migrated over longer distances from the main channel than HUVECs (N = 3). The migration distances were 735.8 ± 330.8 μm for hMSCs and 472.0 ± 220.2 μm for HUVECs-GFP. Aligned dot plot represents mean and st. dev. (** = p < 0.01, ns = not significant; one-way ANOVA). Box plots represent the interquartile range (25th–75th percentile), medians, and whiskers show the full data range (minimum to maximum; ** = p < 0.01, **** = p < 0.0001; ns = not significant; one-way ANOVA).

Subsequently, a subset of the constructs was maintained under static culture conditions, while the remainder was subjected to perfusion culture in a custom-designed bioreactor system (**Figure S11**).^29^ The constructs were perfused with a micropump for the period of 1 week at constant flow rates starting at 300 μL/min for the period of two days to limit cell detachment, following with increased flow rates of 700 μL/min until the end of perfusion. This flow rates resulted in the shear stresses along the channel wall ranging from 0.8 to 2 dyn/cm^2^.^30^ This range of shear stress was hypothesized to improve stability of the main vessel lining without restriction of angiogenic sprouting.^31,32^

Within the macroporous constructs the cells grew into the bulk of the material within the first week of pre-culture; the cells were mostly allocated within the channel in the nanoporous constructs showing limited outgrowth into the striations (**Figure 4c**). The striations that were formed during the printing process create narrow, anisotropic channels and enabled cell invasion into otherwise non-permissive nanoporous materials.^29^ In the macroporous constructs, the hMSCs infiltrated porous space starting from the central channel spanning distances of up to 250 µm within the first 5 days of culture. Interestingly, even though HUVECs-GFP and hMSCs were seeded together they self-organized into an inner layer of endothelial cells (green) surrounded by an outer layer of hMSCs (red) (**Figure 4d**). This arrangement recapitulates the native vascular architecture, where endothelial cells are surrounded by stromal perivascular cells, stabilizing the vessel architecture.^33,34^ HUVECs-GFP had not completely lined the protrusions led by hMSCs but cross sections show partly lumenized endothelial linings in some of the protrusions.

After 2 weeks of culture, macroporous constructs maintained under static and perfused conditions exhibited visibly outgrown vascular structures extending into the bulk of the material. Some of these structures exhibited hierarchical branching (**Figure S12**). The porous constructs maintained under static conditions exhibited cell outgrowth over the distances of 397.4 ± 279.2 μm. The perfused macroporous constructs facilitated evolution of vascular structures spanning over the distances of 520.4 ± 224.0 μm. There was also a limited outgrowth in the nanoporous constructs over the distances of 64.5 ± 29.1 μm and 147.4 ± 83.3, for static and perfused conditions respectively. The outgrowth in nanoporous constructs was mainly allocated within striated regions with locally reduced crosslinking density.^29^ Overall, the macroporosity significantly improved vascularization of the materials bulk compared to nanoporous conditions. While perfusion did not lead to statistically significant improvement in vascular outgrowth under either nano- or macroporous conditions, a trend toward deeper cell infiltration was observed in perfused samples. This suggests that perfusion may enhance outgrowth, though further investigation is needed to confirm this due to large variability observed.

The vessel density was higher in the macroporous compared to nanoporous constructs in both static and perfused conditions. In static conditions the vessel density was 25.3 ± 2.5 mm^-2^ and 61.0 ± 17.1 mm^-2^ for the nano- and macroporous construct respectively. In perfused constructs the vessel density was 21.7 ± 2.8 mm^-2^ and 59.3 ± 5.0 mm^-2^ for the nano- and macroporous construct respectively. Interestingly, while outgrowth distance seemed to be affected by perfusion, vessel density remained largely unaffected. This could be due to the fact that the vessel density is usually regulated through angiogenic growth factors (e.g., VEGF) signalling rather than by the shear forces initiated by perfusion.^35,36^

Within the perfused macroporous chip the migration over the distances of 735.8 ± 330.8 μm for hMSCs and 472.0 ± 220.2 μm for HUVECs-GFP was observed, with hMSCs migrating over longer distances from the main channel than HUVECs-GFP. The longer distances (> 500 μm) were mainly reached through the migration of hMSCs rather than vascular outgrowth. This is likely due to a higher migratory capacity of the hMSCs and their ability to individually navigate porous space, whereas HUVECs typically rely on collective sprouting or need strong angiogenic cues for migration.

## Conclusion

With vascularization being a major challenge in the biofabrication of viable and functional living tissues, there is a clear need for materials that support dynamic cellular processes such as spreading, migration, and tissue infiltration. In this study, we demonstrated that phase-separating photopolymerizable bioinks, when combined with volumetric 3D bioprinting, enable the fabrication of constructs with controllable features on multiple scales – large millimeter-scale porosity derived from the printing process and micrometer-scale porosity arising from phase separation. Furthermore, the use of volumetric light fields not only allows for rapid fabrication but also offers precise control over the extent of phase separation and resulting porosity through modulation of light intensity enabling formation of porosity gradients within the printed structures. This hierarchical pore architecture is particularly well-suited for promoting vascularization in large (centimeter-scale) free-form bioprinted structures. Importantly, the formation of interconnected porous networks addresses the inherent limitations of synthetic hydrogels in supporting cell infiltration and tissue integration. This synergy between light-controlled fabrication and material behavior presents a powerful strategy for engineering functional vascularized tissues.

We generated perfusable constructs with intrinsic porosities that enabled infiltration of vascular structures deep into the bulk of hydrogels. Interestingly, the hMSCs migrated over longer distances than HUVECs. Especially over the long distances (>500 μm), it was predominantly hMSC migration rather than outgrowth of the endothelial structures. These results highlight the need for additional guidance cues, such as gradients of angiogenic factors or mechanical stimuli (e.g., peristaltic flow), to enhance directed vascular outgrowth. Importantly, the VBP platform is highly compatible with the integration of such spatiotemporal biochemical and physical signals, offering a versatile strategy for engineering functional, vascularized tissue and organ-on-chip systems. This approach is well-suited for generating perfusable, macroporous tissue- or organ-on-chip platforms potentially combining them with organoids or spheroids of the target tissue.

## Acknowledgements

This project received funding from the EU under the European Union’s Horizon 2020 research and innovation programme (Grant agreement ID: 964497, ENLIGHT) and from the European Research Council (ERC) under the European Union’s Horizon 2020 research and innovation program (grant agreement No. 949806, VOLUME-BIO). R.L. acknowledges the funding from the Gravitation Program “Materials Driven Regeneration”, funded by the Netherlands Organization for Scientific Research (024.003.013). O.D. acknowledges this publication as a part of the project with file number OCENW.XS22.3.054 of the research programme NWO XS which is (partly) financed by the Dutch Research Council (NWO). We acknowledge C. Labouesse for the help on writing the manuscript, and Paul Delrot (Readily3D) for technical assistance in printing of gradients.

## Author contributions

The project was conceived and designed by O.Y.D., R.L., and M.-B.B., and O.Y.D., R.L., M.-B.B., and G.G. designed experiments. O.Y.D., M.-B.B., G.G., S.A., S.F., A.R.F. carried out the experiments and analyzed data. O.Y.D., R.L. M.-B.B., G.G., M.W.T. wrote the manuscript. All authors have approved the final version of the manuscript.

## Data availability statement

All information necessary to reproduce the work has been includes in the experimental section of the paper. Additional data will be made available upon reasonable request.

## Experimental Section

### Hydrogel production

Hydrogel forming solution for making nanoporous gels were obtained by mixing the following components at final concentrations of 3% w/v GelNb (BioInx), 0.05% w/v LAP (Lithium phenyl(2,4,6-trimethylbenzoyl)phosphinate, Tokyo Chemical Industry, Japan), 0.75 % w/v PEG-SH. The hydrogel mix was always made on the same day as it was used and components were weighed in sterile conditions or sterile filtered for use with cells. The mix was kept in the dark at 37°C and vortexed for at least 10 seconds before being used.

Hydrogel forming solutions for making macroporous gels were obtained by mixing the following components at final concentrations of 3% w/v GelNb (BioInx), 0.05% w/v LAP (Lithium phenyl(2,4,6-trimethylbenzoyl)phosphinate, Tokyo Chemical Industry, Japan), 0.75 % w/v PEG-SH, 0.1 % w/v HA, 3 % w/v dextran. The hydrogel mix was always made on the same day as it was used and components were weighed in sterile conditions or sterile filtered for use with cells. The mix was kept in the dark at 37°C and vortexed for at least 10 seconds before being used.

### Cell culture

hMSCs were isolated from iliac crest bone marrow aspirates obtained from consenting patients (one donor, female; approved by the research ethics committee of the University Medical Center Utrecht, isolation 08-001, distribution protocol 18-739) were cultured in MSC expansion medium (10% FCS, 1% pen/strep, 1% asap in alpha-MEM (Gibco)) supplemented with 5 ng/mL bFGF (PeproTech) on 0.1% w/v gelatin coated cell culture flasks and medium was refreshed twice a week. MSCs were split at a ratio of 1:3 once a week.

GFP-HUVECs were cultured in Endothelial cell growth medium-2 (EGM-2 MV Microvascular Endothelial Cell Growth Medium-2 BulletKit; Lonza; CC-3202) on 0.1 % w/v gelatin coated cell culture flasks and medium was refreshed twice a week. HUVECs were split at a ratio of 1:3 once a week. For cell experiments, passages between 3 to 9 and 6 to 10, for hMSCs and HUVECs-GFP, respectively, were used in this study.

### Light intensity versus porosity

15 µL droplets of hydrogel mix were pipetted onto a PDMS surface. Droplets were exposed to 2.5, 5.6 or 10 mW light at 405 nm for 5 min. Droplets were transferred into a well plate well containing 8 µL/mL Cy3.5 in PBS0 and stored at 4°C overnight to take up the dye. The next day, samples were washed with PBS0 and submerged in dextran-FITC solution for at least 1 hour before imaging using the Leica Stellaris confocal microscope. Z-stacks of 50 µm size were acquired to assess homogeneity of pores throughout the gel.

### Analysis of porosity and pore sizes

The 3D micrograph image stack was processed within ImageJ. First binary thresholding was performed to segment the regions corresponding to the voids (pores) and gel. Porosity was determined by dividing the void volume by the total volume of the imaged sample. To determine the 3D pore sizes, the image stacks were further processed using the plugin BoneJ, enabling skeletonization of the voids, and subsequent determination of pore size distributions within the samples.^24,25^

### Cell seeding or encapsulation in constructs printed via volumetric bioprinting

For volumetric printing, hydrogel mix was pipetted into 10 mm glass vials, vortexed and kept at 37 °C before printing. Additionally, nanoporous conditions were prepared by excluding dextran and HA from the hydrogel mix. Printing was performed with a Tomolite 1.0 bioprinter (Readily3D) and Apparite software. A refractive index of 1.37 and a peak-to-average ratio of 4:1 were set, and printing occurred with 6 mW/cm2 average intensity and 55-60 mJ/cm2 light dose. Post printing, hydrogels were washed thoroughly with warm PBS0. To seed the hydrogels, a cell suspension of 15 uL with a 1:1 hMSCs:GFP-HUVECs ratio for a total of 1 million cells was pipetted on top of the hydrogels. For cell encapsulation studies, a cell pellet containing a 1:1 hMSCs:GFP-HUVECs ratio at 0.5 million cells per mL was resuspended in the hydrogel mix prior to printing, and, post printing, the hydrogels were washed with warm medium. Live/dead staining with the viability/cytotoxicity dyes calcein-AM and ethidium homodimer-1 was performed on days 1 and 3 after printing. The medium was changed every 2 to 3 days, and the hydrogels were imaged with a Thunder Leica microscope.

### Volumetric bioprinting, cell seeding and perfusion of vessel on-a-chip models

Volumetric printing of vessel on-a-chip constructs was executed as previously described.^29,37,38^ Briefly, 1 mL of 37°C hydrogel mix was filled into 10 mm glass vials. Chip models were loaded into apparite software in vertical orientation. Chips were printed at a light dose of 50-70 mJ/cm^2. Chips were washed thoroughly and channels flushed with warm PBS0 and each channel was controlled for perfusability by injecting warm PBS0 into the channel. Chips were glued to well plate wells using 10 % w/v PEGDA with 0.1 % w/v LAP and seeded with MSCs and HUVECs at a 1:1 ratio for a total of 1 million cells. This cell suspension was done in a 7.5 % w/v gelatin solution in medium as the increased viscosity facilitates seeding. After cell injection chips were rotated by 90 degrees along their longitudinal axis every 10 minutes for 6 hours using a custom-built rotating device. Warm 1:1 hMSC:EGM-2 medium was added to the site but not the top of the chips to prevent flushing out the cells. The seeding was repeated twice over a one-week pre-culture period. Then, chips were transferred into a custom-designed perfusion bioreactor system and perfusion was initiated at 300 µL/min for 2 days, and at 700 µL/min until the end of culture. Shear stress, dependent on channel diameter (D), flow rate (Q) and fluid viscosity (η), was calculated according to Equation 1^28^. With channel diameter (D) ≈ 850 μm, flow rate (Q) = [300-700] μL/min, and assuming approximately (η) ≈ 0.001 Pa·s for cell medium viscosity at physiological conditions, these flow rates correspond to shear stress values of 0.8 and 1.9 dyn/cm^2^, respectively.

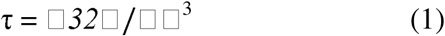

### Volumetric bioprinting of hierarchical porosities

Three cylinders of the same size were inserted into Apparite software and stacked on top of each other. Cylinders were printed in sequential sessions using light intensities of 35 mJ/cm^2^ and 350 mJ/cm^2^ respectively. Cylinders were washed with warm PBS0 and cut into coronal cross sections to assess hierarchical porosities under the bright-field microscope.

### Sample preparation, immunofluorescence stainings and confocal imaging

Samples were prepared as previously described ^34^. Briefly, samples were fixed for 1.5 h at 4°C in 4 % w/v ice cold PFA and subsequently washed with PBT (PBS + 0.1 % w/v Tween20) for at least 30 minutes (or stored in PBT until further processing). Using a vibratome, samples were cut into 350 µm slices. Samples were washed in wash buffer 2 (WB2, 1 L PBS + 1 mL of Triton X-100 + 2 mL of 10% (w/v) SDS + 2 g BSA) for 30 min at 4°C. Samples were stained for CD31 (1:200; Biolegend; 102513), F-actin (1:200; SigmaAldrich 79286-10NMOL) and DAPI (1:1000; Thermo Fisher Scientific Invitrogen™ D1306) overnight at 4°C. Subsequently, samples were washed three times for 1.5 hours in WB2, mounted onto thin glass slides and imaged using the Leica Stellaris or SP8 confocal microscope using a 10, 20 and 25x water immersion objectives.

### Statistical analysis

The experiments were designed and performed with at least three replicates for each condition, and statistical analysis was conducted using Prism 10 (GraphPad). Conditions were assessed using one-way analysis of variance (ANOVA), with Šídák’s multiple comparisons test. *P* values less than 0.05 were considered significant.

---

**Figure S1.**
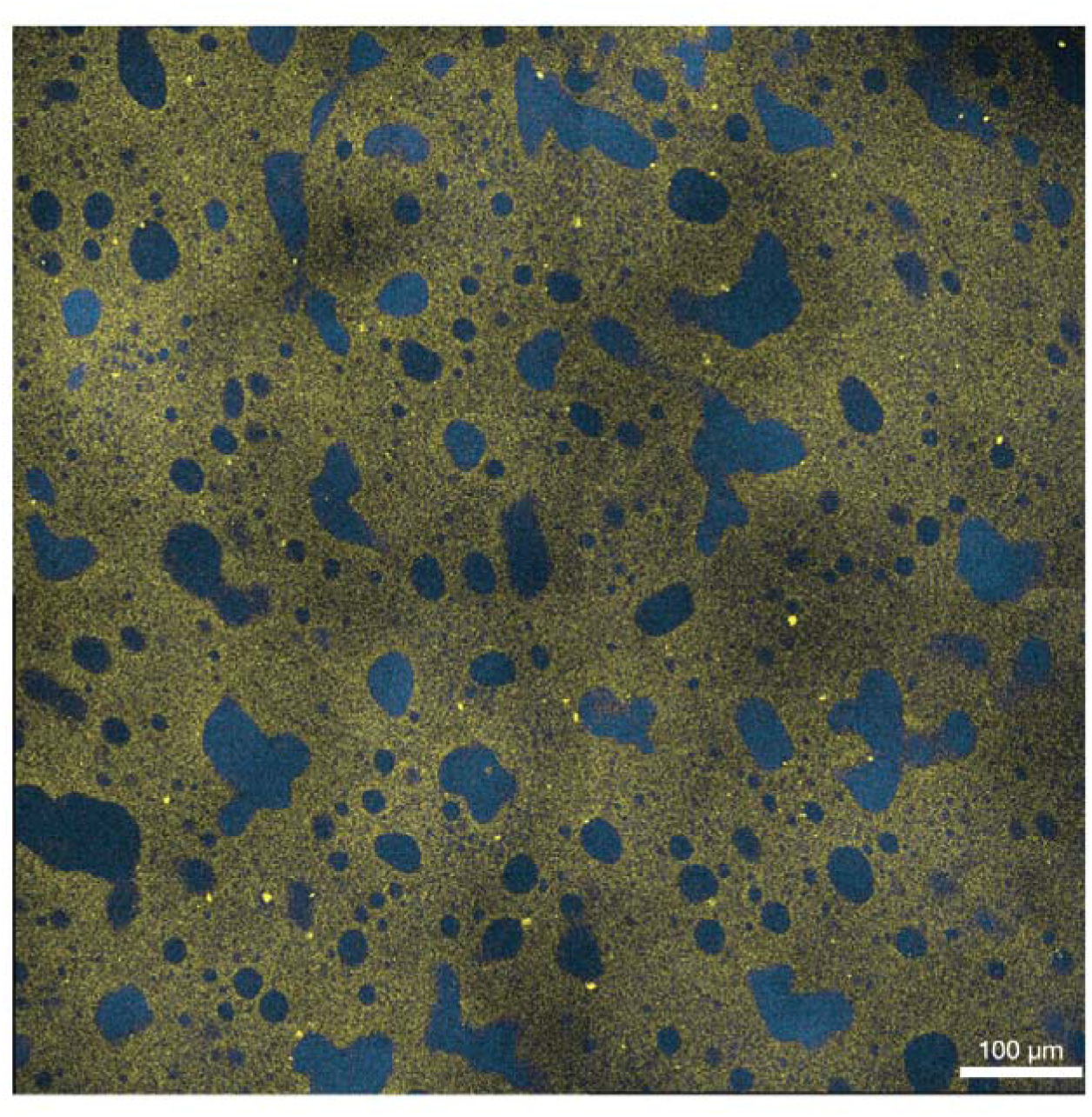
Irregular porosity in GelNb–PEG–SH hydrogels without temperature control. In the absence of temperature regulation during hydrogel cross linking, the GelNb–PEG–SH system forms irregular, binodal structures instead of the desired bicontinuous porosity. The gel phase appears in yellow, while the pore space is visualized in blue using high-molecular-weight Dextran-FITC (500LJkDa).

**Figure S2.**
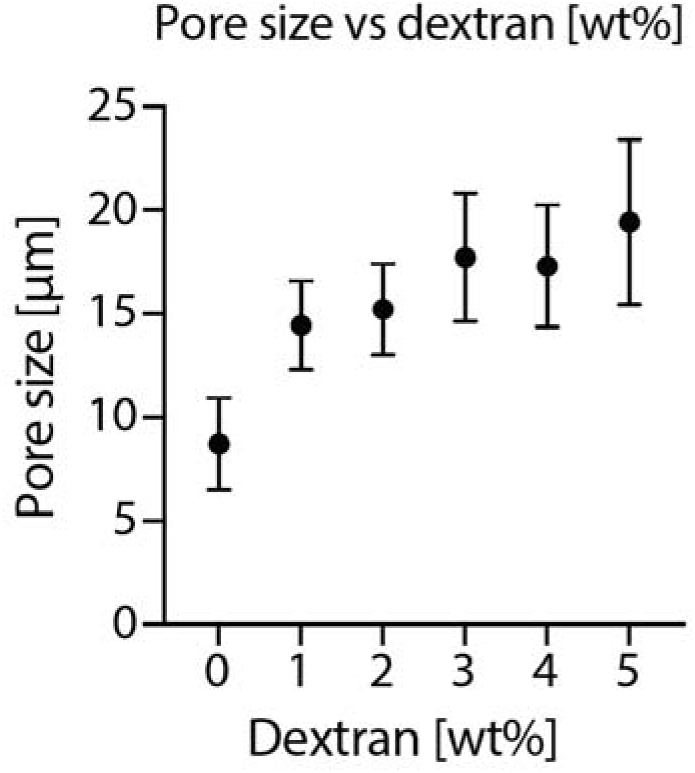
Pore size in 3LJwt% Gel-Nb–PEG–SH hydrogels polymerized under constant light intensit (ILJ=LJ5LJmWLJcm²; nLJ=LJ3) increased at increasing dextran concentration (form 0 to 5LJwt%) . Experimental values are presented as meanLJ±LJstandard deviation (SD).

**Figure S3.**
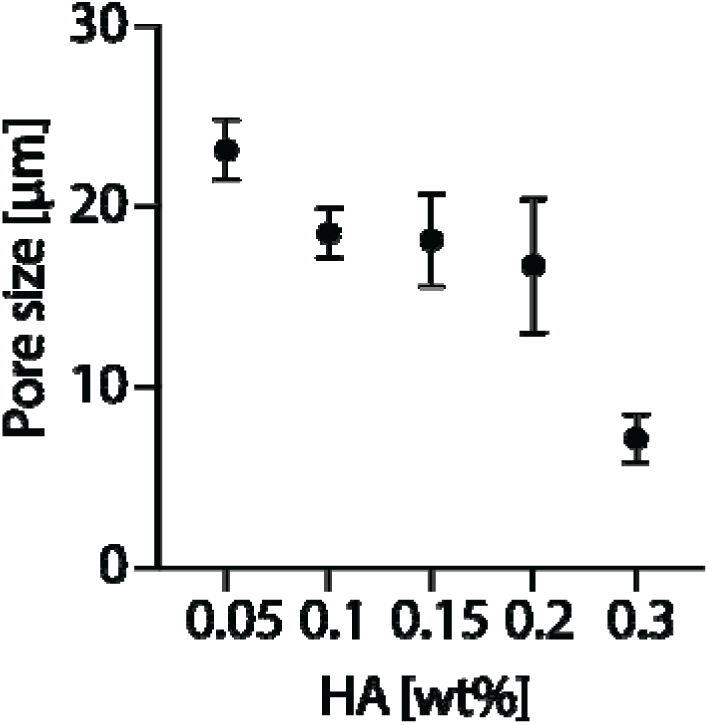
Pore size in 3LJwt% Gel-Nb–PEG–SH hydrogels polymerized under constant light intensit (ILJ=LJ5LJmWLJcm²; nLJ=LJ3) decreased at increasing HA (2 MDa) concentration (from 0.05 to 0.3LJwt%) . Experimental values are presented as meanLJ±LJstandard deviation (SD).

**Figure S4.**
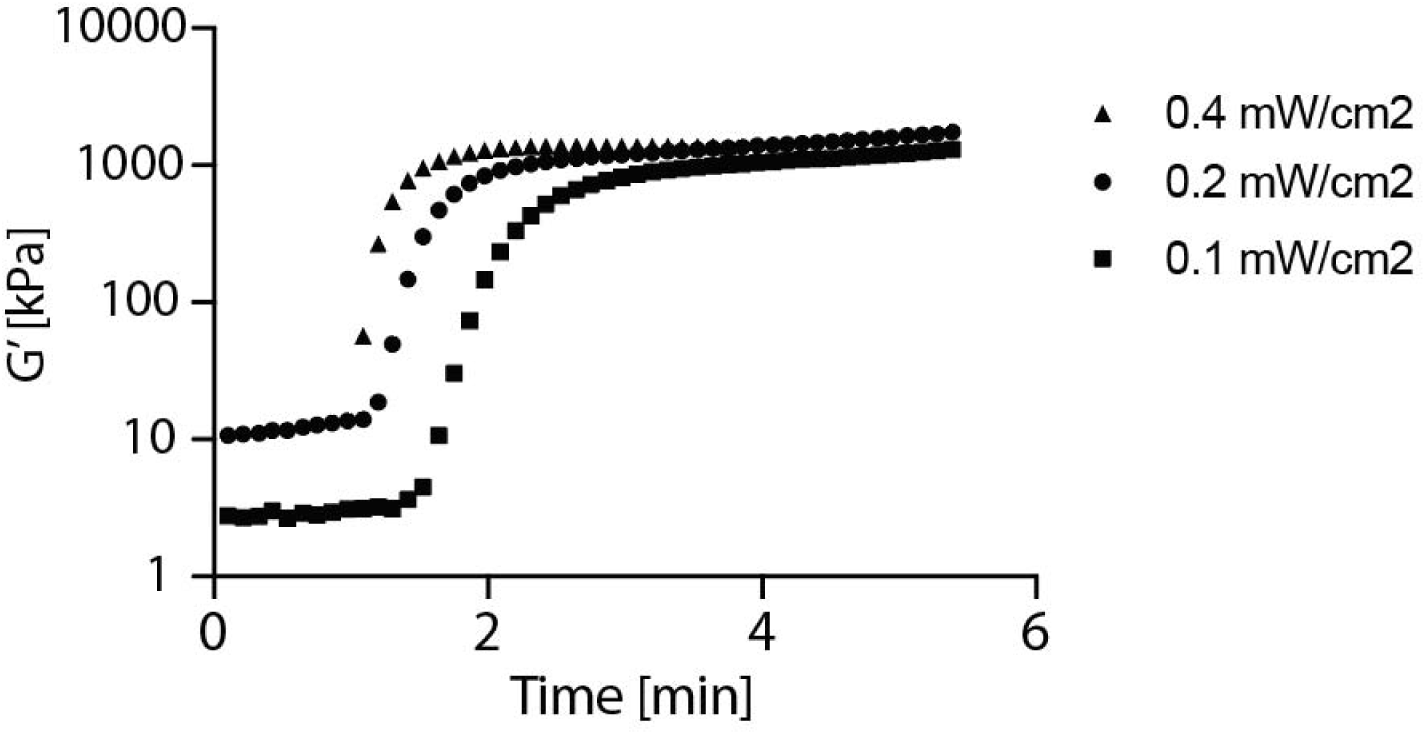
Influence of irradiation intensity on gelation kinetics in Gel-NB–PEG–SH hydrogels. The evolution of the storage modulus (G’) during photopolymerization was monitored using a rheometer to assess gelation kinetics in Gel-NB–PEG–SH hydrogels (3 wt%) exposed to varying light intensities. All samples were irradiated starting at the same time point (1 minute after measurement onset). Lower light intensities led to slower increases in G’ and delayed plateau formation, indicating that crosslinking kinetics are strongly dependent on irradiation intensity. The hydrogels crosslinked at different light intensities reacht the same plateau modulus.

**Figure S5.**
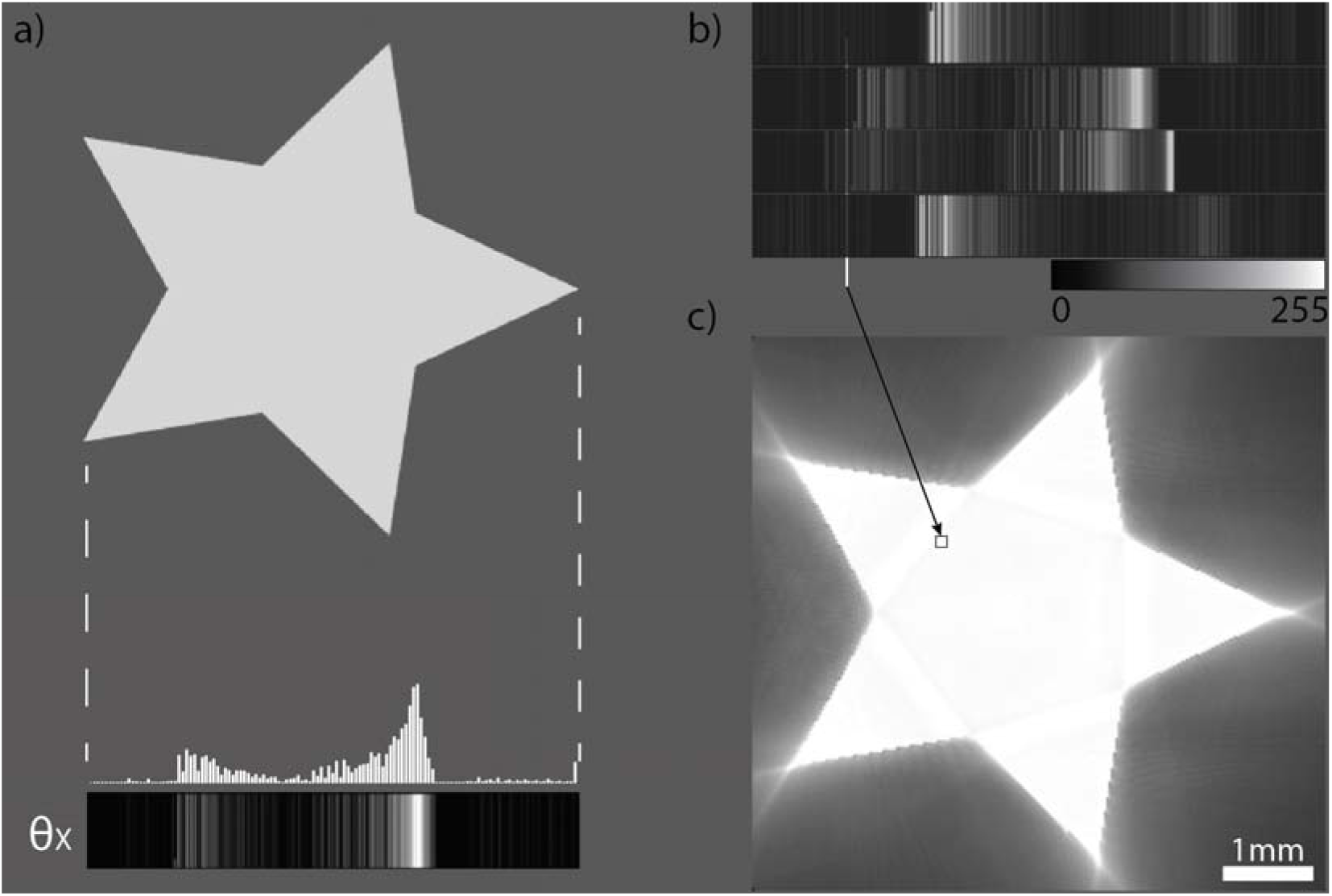
Intensity distribution during volumetric printing. a) An example of a projected pattern at an arbitrary angle (θx) for the tomographic projection of a star, demonstrating the non-homogeneous intensity distribution at each projected angle. b) Four example projections each generated at a different projection angle, in which a 2D pixel is targeting the same voxel. While the projected pixels vary in light intensity at each projected angle, the sum of all pixels reaches the photocrosslinking threshold within the object’s voxel (arrow targets a hypothetical voxel volume). In contrast, absolute grayscale values are linked to an absolute light intensity. c) The filtered tomographic backprojection of the printed star, showing the final reconstructed light dose distribution.

**Figure S6.**
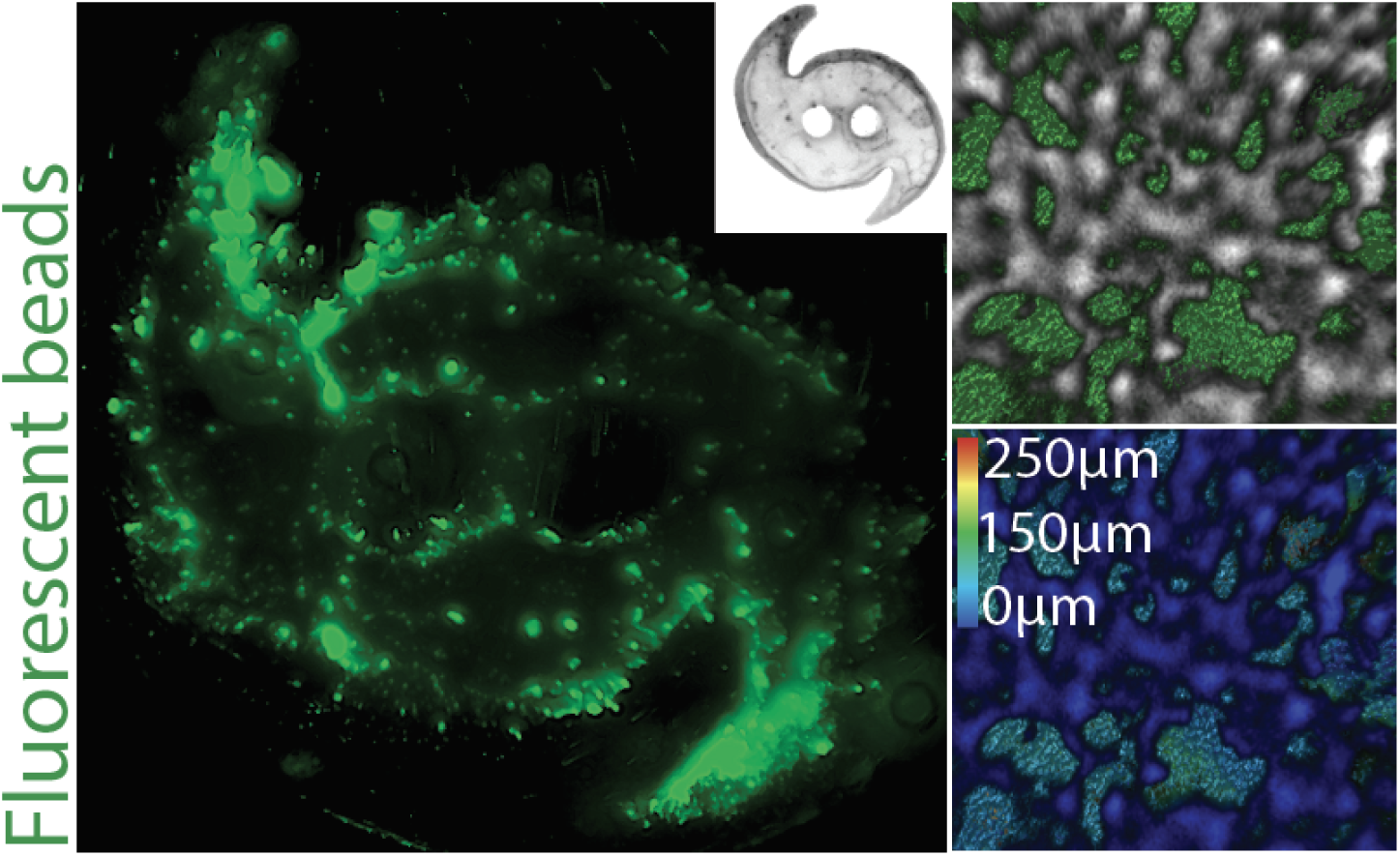
Maximum projection of the macroporous galaxy print that was agitated with the green fluorescent microbeads (d = 6 μm) with the insert of the corresponding printed shape. The interconnected porous space allowed the penetration of microbeads into the construct (left). The 3D reconstruction of the porous space shows the accumulation of the microbeads (green) within the pores of the hydrogel (gray) to the depths of 150 μm.

**Figure S7.**
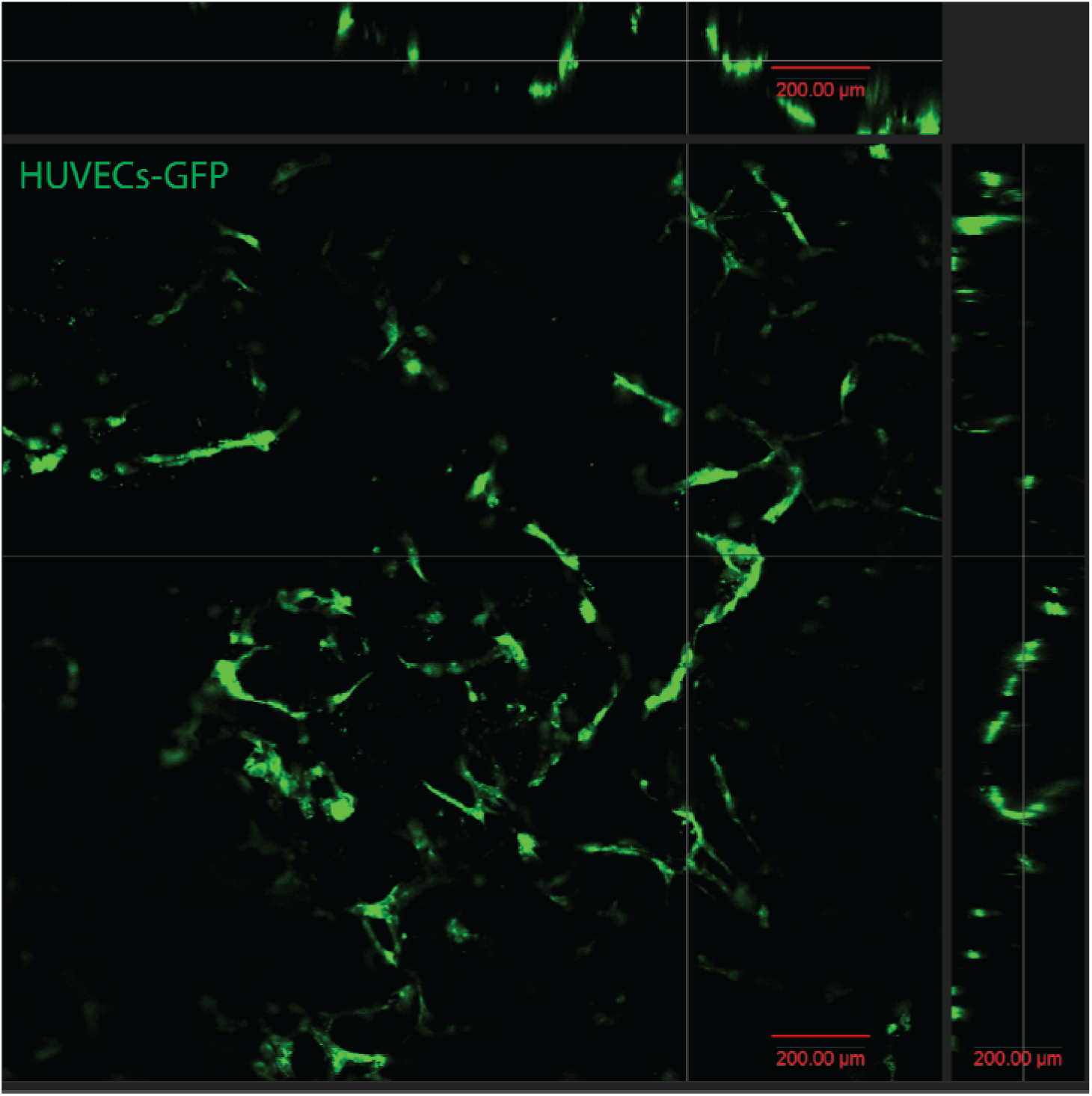
Infiltration of HUVECs-GFP into porous hydrogels. HUVECs-GFP were seeded onto the surface of porous hydrogel constructs. Wishing 5 day of the culture period, the cells actively migrated and infiltrated into the pore space, reaching depths of up to 300LJμm. Imaging was performed using confocal microscopy to visualize and quantify the depth of cellular penetration within the scaffold.

**Figure S8.**
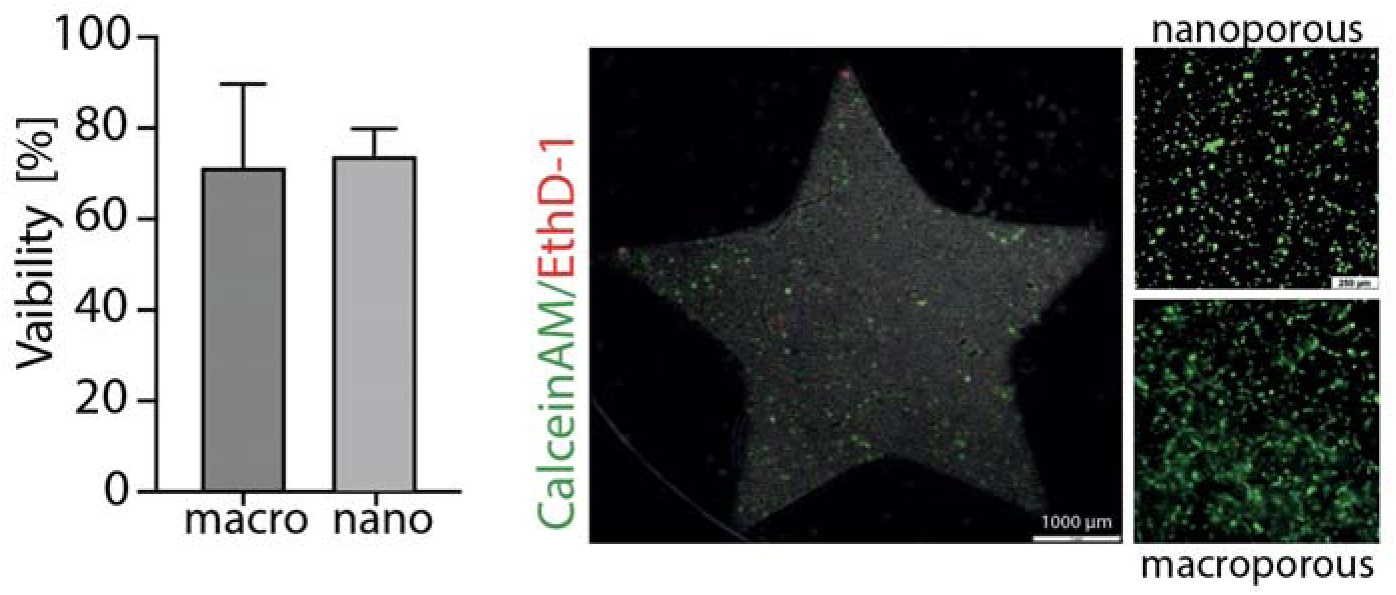
Viability of printed cells within star-shaped constructs. The viability of HUVECs-GFP and hMSCs encapsulated in star-shaped constructs printed under varying light intensities (4–10LJmW/cm²) was assessed 24 hours post-printing. Cell viability was quantified using live/dead (Calcein-AM/Ethidium Homodimer-1) staining and fluorescence imaging. In nanoporous hydrogels, viability was 73.8LJ±LJ5.7%, while in macroporous hydrogels it was 71.3LJ±LJ18.0%, indicating comparable post-printing cell survival across both hydrogel architectures.

**Figure S9.**
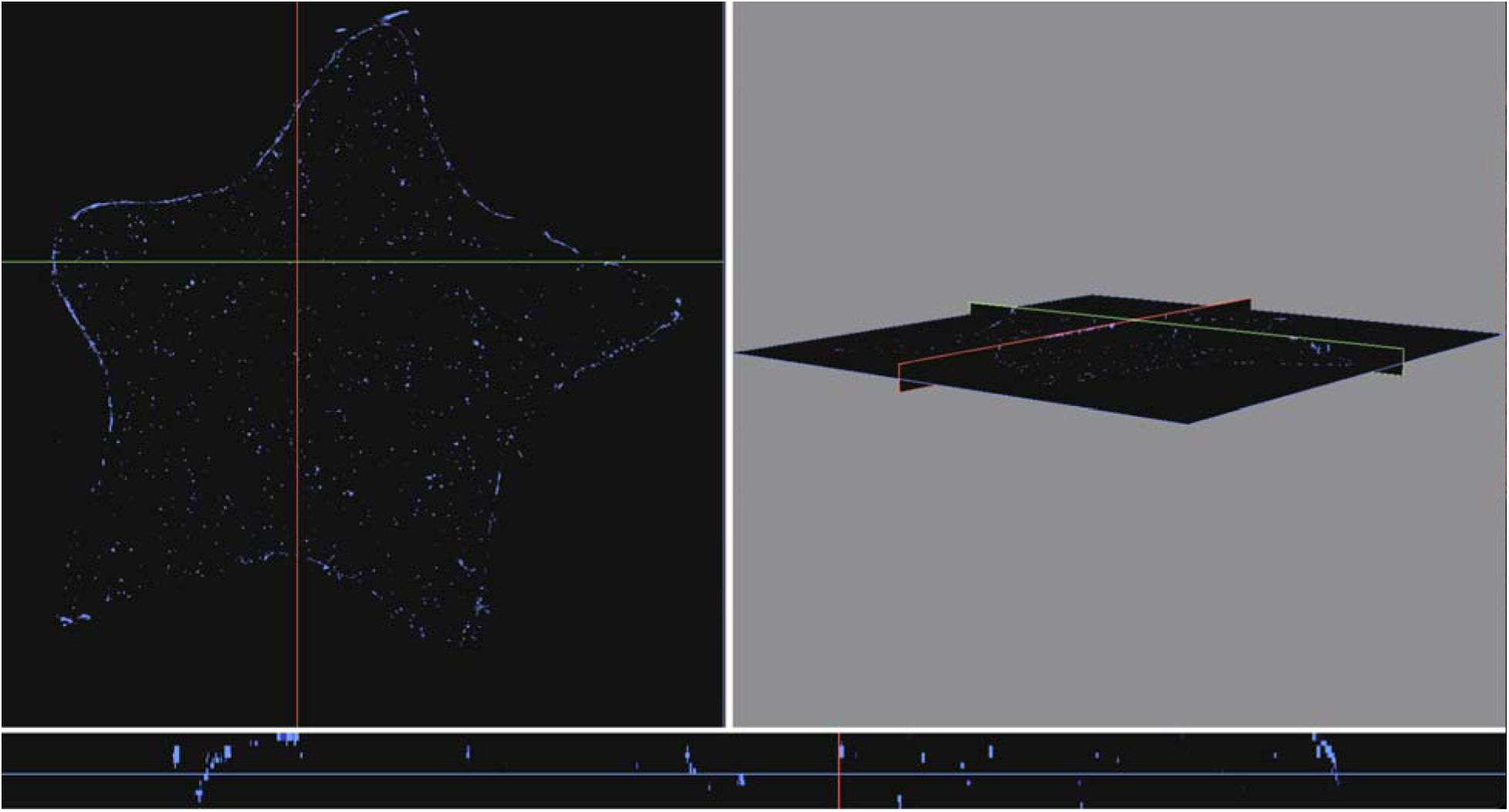
The distribution of DAPI signals within the volume of printed star constructs reconstructed from the z-stack of the confocal slices (total z-dimension length is 50 μm).

**Figure S10.**
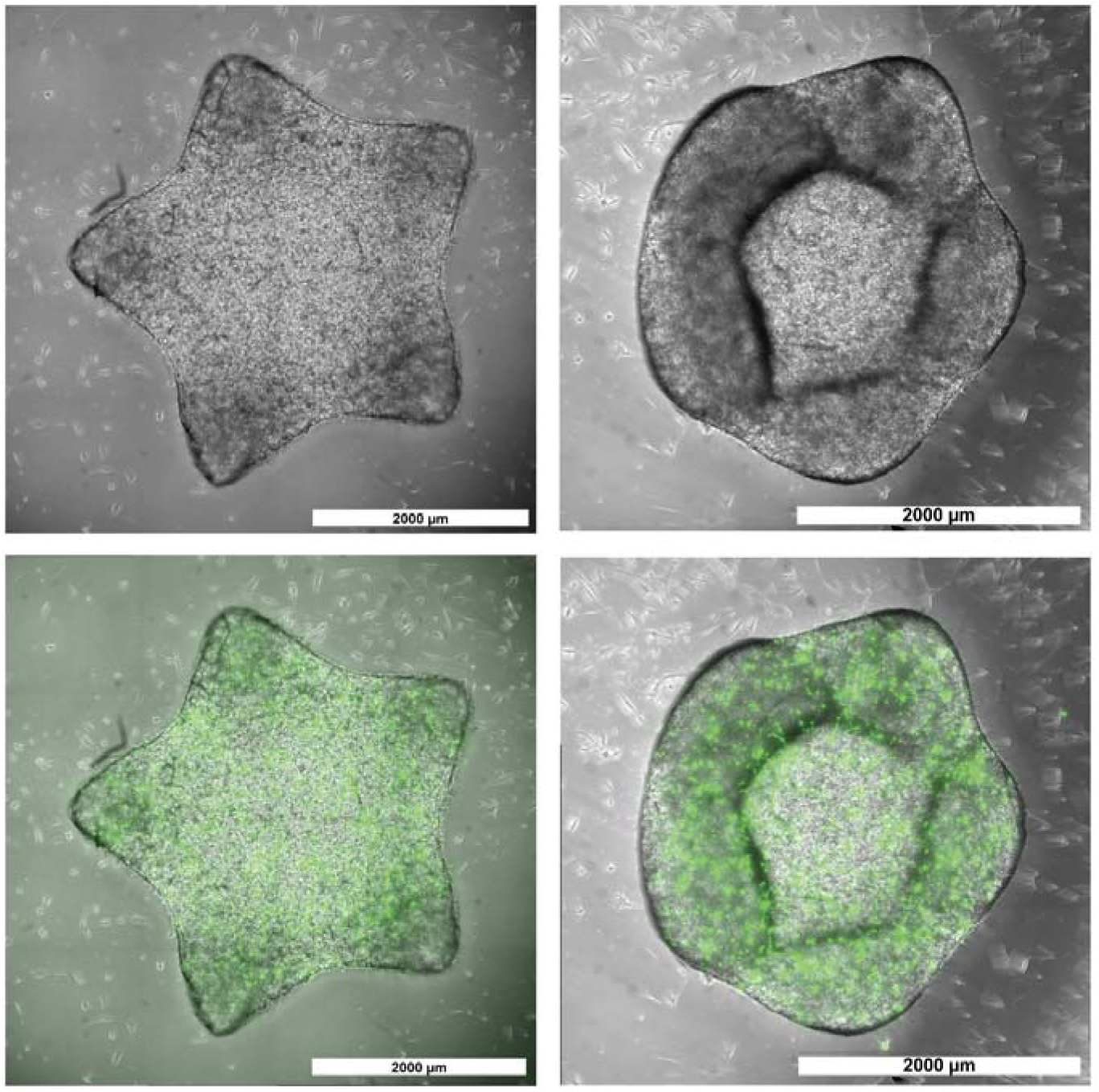
Shrinking of macroporous star-shaped constructs. Macroporous star-shaped constructs with a thickness of 0.5LJmm supported the formation of endothelial structures throughout the hydrogel matrix. Over a 7-day culture period, some constructs exhibited noticeable shrinkage. Despite this macroscopic contraction, the internal endothelial networks remained structurally stable.

**Figure S11.**
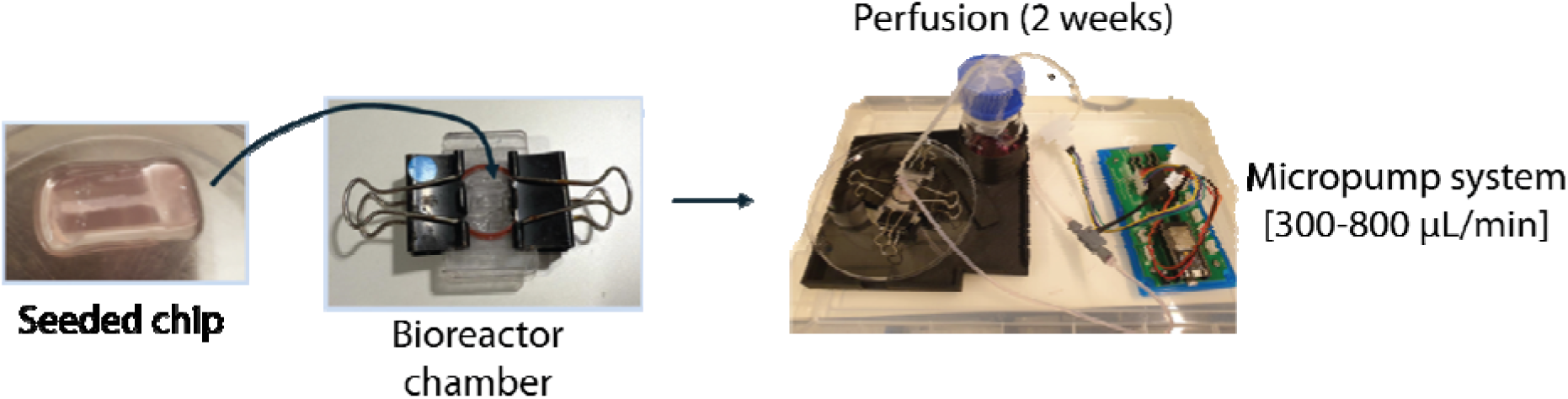
Perfusion set-up. The printed constructs that were seeded with HUVECs-GFP and hMSCs were placed in a bioreactor chamber equipped for connection to the perfusion tubing. The chamber was then connected to a micropump perfusion system, which continuously perfused the chips with fresh medium (EGM-2/alpha-MEM) to support HUVECs and hMSCs over a 2-week period under physiological conditions.

**Figure S12.**
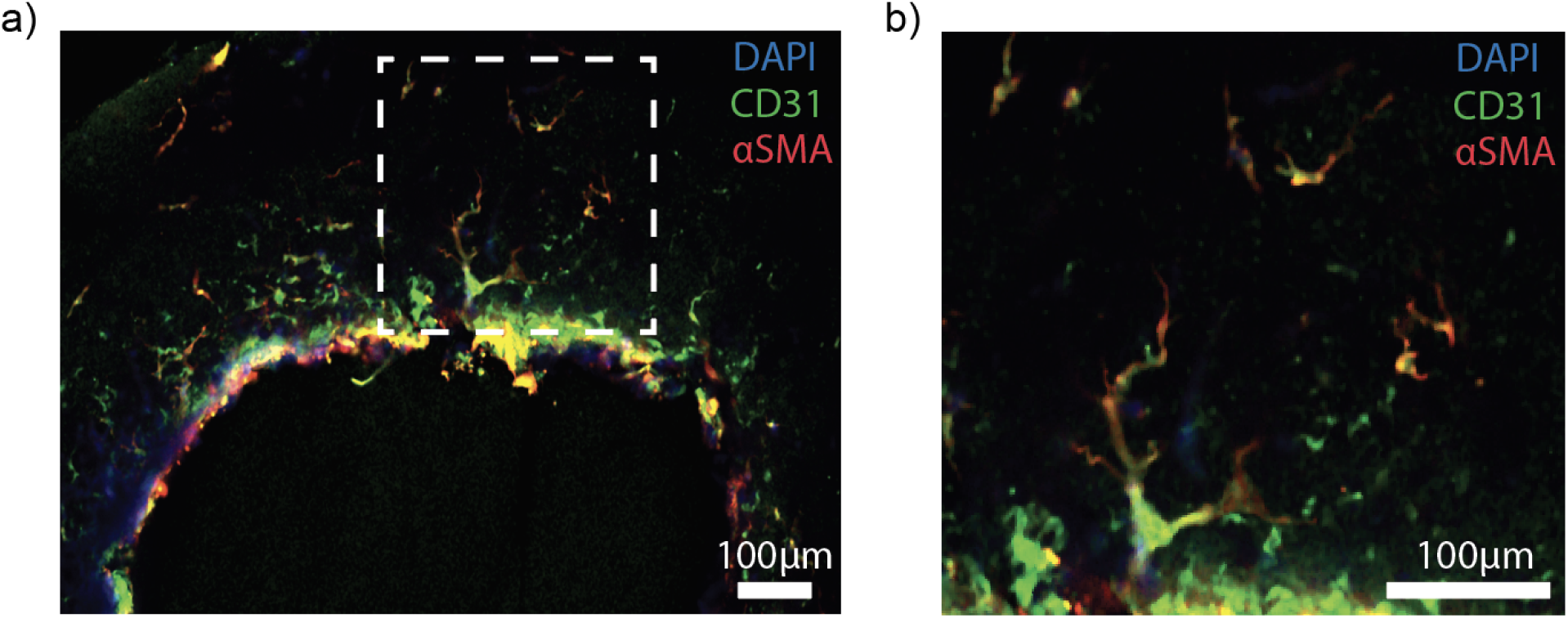
Hierarchical vascular branching from the main perfused channel. A radial cross-section of the main channel in the perfusable construct after 2 weeks of perfusion culture reveals the formation of organized, hierarchical vascular branches extending from the main channel into the surrounding porous hydrogel matrix.

## References

1. Dai, D. et al. Multimorbidity in Atherosclerotic Cardiovascular Disease and Its Associations With Adverse Cardiovascular Events and Healthcare Costs: A Real-World Evidence Study. Journal of Health Economics and Outcomes Research 11, 75 (2024).

2. Recent advances in biofabrication strategies based on bioprinting for vascularized tissue repair and regeneration. Materials & Design 229, 111885 (2023).

3. Kaully, T., Kaufman-Francis, K., Lesman, A. & Levenberg, S. Vascularization--the conduit to viable engineered tissues. Tissue Eng Part B Rev 15, 159–169 (2009).

4. Polacheck, W. J., Kutys, M. L., Tefft, J. B. & Chen, C. S. Microfabricated blood vessels for modeling the vascular transport barrier. Nature Protocols 14, 1425–1454 (2019).

5. Sundaram, S. et al. Sacrificial capillary pumps to engineer multiscalar biological forms. Nature 636, 361–367 (2024).

6. Bott, K. et al. The effect of matrix characteristics on fibroblast proliferation in 3D gels. Biomaterials 31, 8454–8464 (2010).

7. Wei, Z. et al. Hydrogels with tunable mechanical plasticity regulate endothelial cell outgrowth in vasculogenesis and angiogenesis. Nature Communications 14, 1–16 (2023).

8. Bernal, P.N. et al., Volumetric Bioprinting of Complex Living-Tissue Constructs within Seconds. Advanced Materials 31,e1904209 (2019).

9. Bernal, P.N. et al., The road ahead in materials and technologies for volumetric 3D printing. Nature Reviews Materials (2025). doi: 10.1038/s41578-025-00785-3

10. Datta, P., Ayan, B. & Ozbolat, I. T. Bioprinting for vascular and vascularized tissue biofabrication. Acta Biomater 51, 1–20 (2017).

11. Grigoryan, B. et al. Multivascular networks and functional intravascular topologies within biocompatible hydrogels. Science 364, 458–464 (2019).

12. Levato, R. et al. High-resolution lithographic biofabrication of hydrogels with complex microchannels from low-temperature-soluble gelatin bioresins. Materials Today Bio 12, 100162 (2021).

13. Bernal, P. N. et al. Volumetric Bioprinting of Organoids and Optically Tuned Hydrogels to Build Liver-Like Metabolic Biofactories. Adv Mater 34, e2110054 (2022).

14. Dobos, A. et al. On-chip high-definition bioprinting of microvascular structures. Biofabrication 13, 015016 (2021).

15. Cantoni, F., Barbe, L., Pohlit, H. & Tenje, M. A Perfusable Multi-Hydrogel Vasculature On-Chip Engineered by 2-Photon 3D Printing and Scaffold Molding to Improve Microfabrication Fidelity in Hydrogels. Advanced Materials Technologies 9, 2300718 (2024).

16. Rayner, S. G. et al. Multiphoton-Guided Creation of Complex Organ-Specific Microvasculature. Adv Healthc Mater 10, e2100031 (2021).

17. Lévesque, S. G., Lim, R. M. & Shoichet, M. S. Macroporous interconnected dextran scaffolds of controlled porosity for tissue-engineering applications. Biomaterials 26, 7436– 7446 (2005).

18. Broguiere, N. et al. Macroporous hydrogels derived from aqueous dynamic phase separation. Biomaterials 200, 56–65 (2019).

19. Dudaryeva, O. Y. et al. Tunable Bicontinuous Macroporous Cell Culture Scaffolds via Kinetically Controlled Phase Separation. Adv Mater 37, e2410452 (2025).

20. Ying, G.-L. et al. Aqueous Two-Phase Emulsion Bioink-Enabled 3D Bioprinting of Porous Hydrogels. Adv Mater 30, e1805460 (2018).

21. Gonella, S., et al. Fabrication and Characterization of Porous PEGDA Hydrogels for Articular Cartilage Regeneration. Gels 10, (2024).

22. Zauchner, D. et al. Synthetic biodegradable microporous hydrogels for in vitro 3D culture of functional human bone cell networks. Nat Commun 15, 5027 (2024).

23. Müller, M. Z. et al. Cell-guiding microporous hydrogels by photopolymerization-induced phase separation. Nat Commun 16, 4923 (2025).

24. Doube, M. et al. BoneJ: Free and extensible bone image analysis in ImageJ. Bone 47, 1076–1079 (2010).

25. Domander, R., Felder, A. A. & Doube, M. BoneJ2 - refactoring established research software. Wellcome Open Res 6, 37 (2021).

26. Guo, D., Wang, Q., Li, C., Wang, Y. & Chen, X. VEGF stimulated the angiogenesis by promoting the mitochondrial functions. Oncotarget 8, 77020–77027 (2017).

27. Nguyen, D.-H. T. et al. Biomimetic model to reconstitute angiogenic sprouting morphogenesis in vitro. Proc Natl Acad Sci U S A 110, 6712–6717 (2013).

28. Galie, P. A. et al. Fluid shear stress threshold regulates angiogenic sprouting. Proceedings of the National Academy of Sciences 111, 7968–7973 (2014).

29. Buchholz, M.-B. et al. Development of a bioreactor and volumetric bioprinting protocol to enable perfused culture of biofabricated human epithelial mammary ducts and endothelial constructs. Biofabrication (2025) doi:10.1088/1758-5090/add20f.

30. Roux, E., Bougaran, P., Dufourcq, P. & Couffinhal, T. Fluid Shear Stress Sensing by the Endothelial Layer. Front Physiol 11, 861 (2020).

31. Lee, V. K. et al. Generation of Multi-Scale Vascular Network System within 3D Hydrogel using 3D Bio-Printing Technology. Cell Mol Bioeng 7, 460–472 (2014).

32. Lee, V. K. et al. Creating perfused functional vascular channels using 3D bio-printing technology. Biomaterials 35, 8092–8102 (2014).

33. Yang, G., Mahadik, B., Choi, J. Y. & Fisher, J. P. Vascularization in tissue engineering: fundamentals and state-of-art. *Progress in biomedical engineering (Bristol*, England) 2, 012002 (2020).

34. Pill, K. et al. Microvascular Networks From Endothelial Cells and Mesenchymal Stromal Cells From Adipose Tissue and Bone Marrow: A Comparison. Front Bioeng Biotechnol 6, 156 (2018).

35. Lund, E. L., Thorsen, C., Pedersen, M. W., Junker, N. & Kristjansen, P. E. Relationship between vessel density and expression of vascular endothelial growth factor and basic fibroblast growth factor in small cell lung cancer in vivo and in vitro. Clin Cancer Res 6, 4287–4291 (2000).

36. Ylä-Herttuala, S., Rissanen, T. T., Vajanto, I. & Hartikainen, J. Vascular endothelial growth factors: biology and current status of clinical applications in cardiovascular medicine. J Am Coll Cardiol 49, 1015–1026 (2007).

37. Ribezzi, D. et al., Multi-material Volumetric Bioprinting and Plug-and-play Suspension Bath Biofabrication via Bioresin Molecular Weight Tuning and via Multiwavelength Alignment Optics. Advanced Materials 37, 2409355 (2025).

38. Falandt, M. et al., Spatial-Selective Volumetric 4D printing and Single-Photon Grafting of Biomolecules within Centimeter-Scale Hydrogels via Tomographic Manufacturing. Advanced Materials Technologies 8, 2300026 (2023).

